# Evolutionary conservation of midline axon guidance activity between *Drosophila* and *Tribolium* Frazzled

**DOI:** 10.1101/2024.12.20.629797

**Authors:** Piyasi Ghosh, Benjamin C. Wadsworth, Logan Terry, Timothy A. Evans

## Abstract

The regulation of midline crossing of axons is of fundamental importance for the proper development of nervous system connectivity in bilaterian animals. A number of conserved axon guidance signaling pathways coordinate to attract or repel axons at the nervous system midline to ensure the proper regulation of midline crossing. The attractive Netrin-Frazzled/DCC (Net-Fra) signaling pathway is widely conserved among bilaterians, but it is not clear whether the mechanisms by which Net and Fra promote midline crossing are also conserved. In *Drosophila*, Fra can promote midline crossing via Netrin-dependent and Netrin-independent mechanisms, by acting as a canonical midline attractive receptor and also through a non-canonical pathway to inhibit midline repulsion via transcriptional regulation. To examine the conservation of Fra-dependent axon guidance mechanisms among insects, in this paper we compare the midline attractive roles of the Frazzled receptor in the fruit fly (*Drosophila melanogaster*) and flour beetle (*Tribolium castaneum*) using CRISPR/Cas9-mediated gene editing. We replace the *Drosophila fra* gene with sequences encoding *Drosophila* Fra (DmFra) or *Tribolium* Fra (TcFra) and examine midline crossing of axons in the ventral nerve cord of embryos carrying these modified alleles. We show that *Tribolium* Fra can fully substitute for *Drosophila* Fra to promote midline crossing of axons in the embryonic nervous system, suggesting that the mechanisms by which Frazzled regulates midline axon guidance may be evolutionarily conserved within insects.

## Introduction

During nervous system development, evolutionarily conserved signaling pathways direct growing axons towards their synaptic targets. Axon guidance typically occurs in stages, as individual axons respond to a series of cues which direct specific guidance decisions at choice points along the axon’s intended trajectory. In the embryonic central nervous system (CNS) of bilaterian animals, most axons must choose whether or not to cross the midline to the opposite side of the body (Evans and Bashaw, 2010). This fundamental axon guidance choice is regulated by attractive signaling pathways such as the Netrin-Frazzled/DCC (Net-Fra) pathway (Chan et al., 1996; Harris et al., 1996; Kennedy et al., 1994; Kolodziej et al., 1996; Mitchell et al., 1996; Serafini et al., 1994, 1996; Cebrià and Newmark, 2005; Linne and Stollewerk, 2011), and repulsive signaling pathways like the Slit-Roundabout (Slit-Robo) pathway (Cebrià et al., 2007; Cebrià and Newmark, 2007; Evans and Bashaw, 2012; Fricke et al., 2001; Kidd et al., 1998; Long et al., 2004; Zallen et al., 1998). In insect embryos, the precise regulation of midline crossing produces a uniformly segmented ladder-like arrangement of axons in the ventral nerve cord neuropile, with commissural axons crossing the midline at two specific locations in each segment (the anterior and posterior commissures; AC and PC) to produce the “rungs” of the ladder (Evans and Bashaw, 2012; Thomas et al., 1984).

Much of our understanding of the molecular and genetic regulation of axon guidance and midline crossing has come from studies in the fruit fly *Drosophila melanogaster*. In *Drosophila*, the Frazzled (Fra) receptor promotes midline crossing through two distinct mechanisms: by signaling midline attraction in response to its canonical Netrin ligands (NetA and NetB, both expressed at the ventral nerve cord midline) (Harris et al., 1996; Kolodziej et al., 1996; Mitchell et al., 1996), and by inhibiting midline repulsion through a non-canonical pathway involving proteolytic processing of Fra and nuclear translocation of its intracellular domain, which activates transcription of *commissureless (comm)*, a negative regulator of the repulsive Slit-Robo pathway (Neuhaus-Follini and Bashaw, 2015; Yang et al., 2009; Zang et al., 2022).

Despite its prominent role in signaling midline attraction, many axons in *Drosophila* still cross the midline in the absence of the Net-Fra pathway (Harris et al., 1996; Kolodziej et al., 1996; Mitchell et al., 1996). Clearly, additional factors or pathways must be able to promote midline crossing when Netrin or Frazzled are disabled. Indeed, a number of genes have been identified in *Drosophila* which strongly enhance midline crossing defects when their functions are removed in *NetAB* or *fra* mutant backgrounds, including *robo2* (Evans et al., 2015; Spitzweck et al., 2010), *semaphorin-1a (sema-1a)* (Hernandez-Fleming et al., 2017), *flamingo (fmi)* (Organisti et al., 2014), and *Down syndrome cell adhesion molecule (Dscam)* (Andrews et al., 2008).

Although the main components of midline attractive and repulsive pathways are evolutionarily conserved, there is evidence that the specific mechanisms by which they regulate midline crossing may be different, even within insects (Evans, 2016). For example, while Slit and its Robo receptors do signal midline repulsion in the flour beetle *Tribolium castaneum, comm* does not appear to be conserved in non-dipteran insects like *Tribolium*, and *Tribolium* Robo2/3 appears unable to promote midline crossing of axons in *Drosophila* embryos, unlike its *Drosophila* ortholog Robo2 (Evans and Bashaw, 2012). In addition, knockdown of *frazzled* in the mosquito *Aedes aegypti* produces a much stronger reduction in midline crossing than seen in *Drosophila fra* or *NetAB* mutants, suggesting that the additional factors that promote midline crossing in *Drosophila* independently of Net and Fra may not function this way in mosquitoes (Clemons et al., 2011).

We have previously compared the roles of the Slit-Robo pathway in axon guidance in *Drosophila* and *Tribolium*, using RNAi knockdown approaches in *Tribolium* combined with transgenic and gene replacement approaches in *Drosophila*, and observed conservation of some features of pathway function and regulation, and divergence of others (Evans, 2017; Evans and Bashaw, 2012). Others have described conserved features of Netrin and Frazzled expression in *Drosophila* and *Tribolium* (Janssen and Budd, 2023; Simanton et al., 2009), but it has not yet been determined whether the Net-Fra pathway promotes midline crossing in *Tribolium*, or whether *Drosophila* and *Tribolium* Frazzled may guide axons via the same mechanisms. Here, we compare the functions of Frazzled in *Tribolium* and *Drosophila* using CRISPR/Cas9-mediated gene replacement. We create engineered alleles of *Drosophila fra* in which the coding region is replaced by cDNAs encoding either *Drosophila* Fra (DmFra) or *Tribolium* Fra (TcFra), and show that the *Tribolium* Fra protein is stably expressed and properly localized in *Drosophila* neurons in vivo, and that TcFra can fully substitute for DmFra to promote midline crossing and longitudinal guidance of axons in the *Drosophila* embryonic CNS. Our results support the hypothesis that the mechanisms by which Fra promotes midline crossing are evolutionarily conserved within insects.

## Results

### Identification of *Tribolium frazzled*

To identify *frazzled* orthologs in *Tribolium*, we searched the *Tribolium castaneum* genome by tblastn using the *Drosophila melanogaster* Frazzled (Fra) protein as a query sequence.

Consistent with a previous analysis by Janssen and Budd (Janssen and Budd, 2023), we identified a single *Tribolium* genomic region encoding a Fra-like protein. We used PCR primers designed against the predicted exons encoding the N-terminal Ig1 domain and stop codon to amplify and clone partial cDNAs from 0-6 day old *Tribolium* embryos. We identified two alternatively spliced isoforms which we designated *TcFraL* (long isoform) and *TcFraS* (short isoform). Both isoforms encode transmembrane proteins containing four extracellular immunoglobulin-like (Ig) domains and six fibronectin (Fn) repeats, similar to *Drosophila* Frazzled and its vertebrate orthologs Deleted in Colorectal Cancer (DCC) and Neogenin (Keino-Masu et al., 1996; Kolodziej et al., 1996; Vielmetter et al., 1994). We hereafter refer to the *Drosophila melanogaster* Fra protein as “DmFra” to distinguish it from *Tribolium castaneum* Fra (“TcFra”).

The two *Tribolium* Frazzled isoforms differ due to alternative splicing at two locations, with the longer isoform (TcFraL) including an additional 98 amino acids between the Ig4 and Fn1 domains and 13 amino acids between Fn6 and the transmembrane domain. Notably, the *Drosophila fra* gene is also alternatively spliced, with the longer Fra isoform including a 151 aa insertion at the same location between Ig4 and Fn6 (Kolodziej et al., 1996). The functional significance of this alternative splicing in *Drosophila fra* is not known, and the extra sequence in the long DmFra and TcFra isoforms do not appear to share significant primary sequence similarity. The three previously identified evolutionarily conserved cytoplasmic sequence motifs in DmFra (P1, P2, P3) are also present in both TcFra isoforms. A schematic of the *Tribolium frazzled* locus and sequence alignment of the DmFra and TcFra proteins are shown in **Figure 1**.

**Figure 1.**
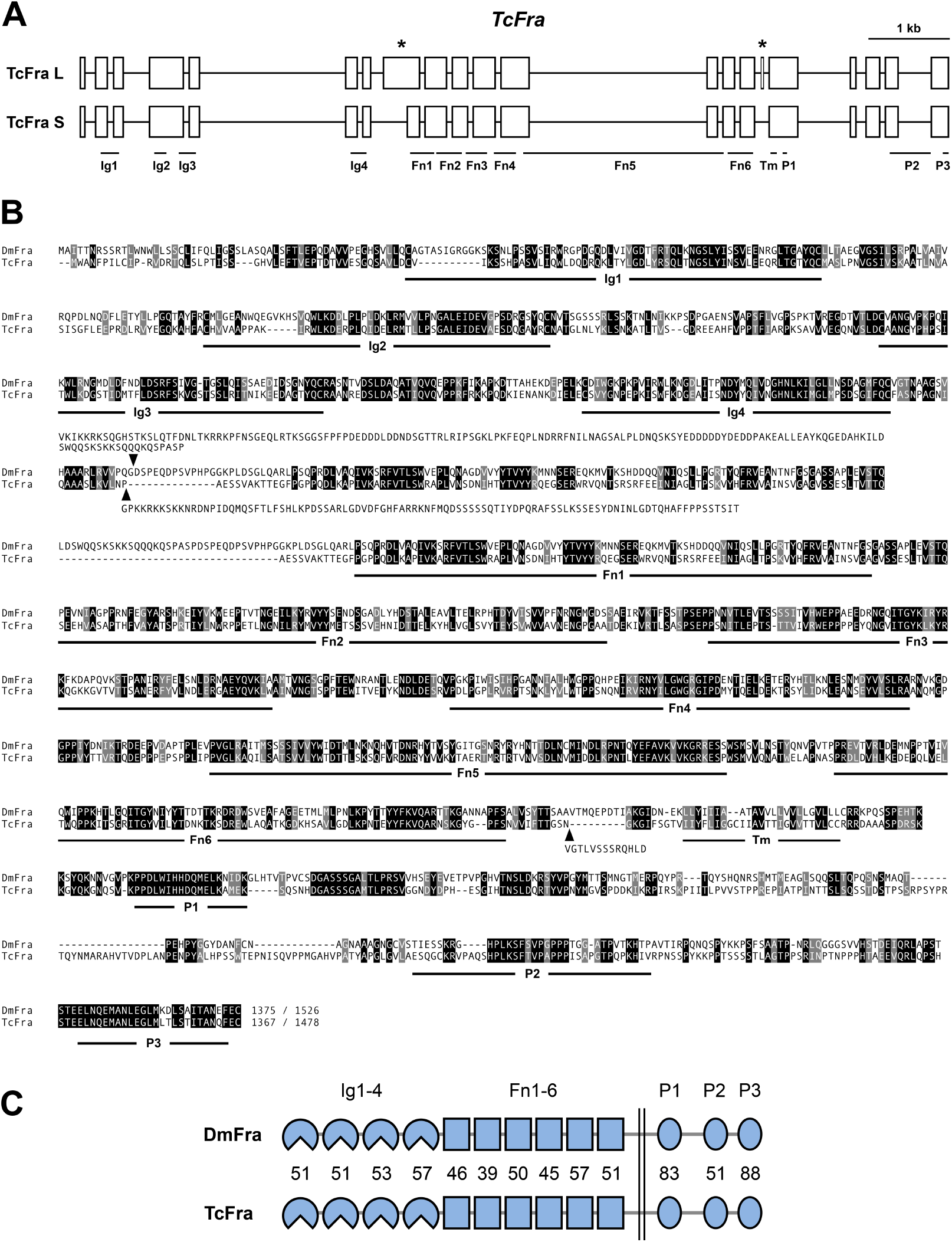
Sequence comparison of *Drosophila* and *Tribolium* Frazzled. (**A**) Schematic of the *Tribolium frazzled (TcFra)* gene. Boxes indicate exons. *TcFra* is alternatively spliced at two locations (asterisks) to produce long *(TcFraL)* and short *(TcFraS)* isoforms. (**B**) Sequence alignment of *Drosophila* Fra (DmFra) and *Tribolium* Fra (TcFra) proteins. Structural features are indicated below the sequence. Identical residues are shaded black; similar residues are shaded gray. Black arrowheads show locations of alternative splicing, and the additional sequence found in the long isoforms is shown above (for DmFra) or below (for TcFra). Ig, immunoglobulin-like domain; Fn, fibronectin repeat; Tm, transmembrane helix; P1-P3, conserved cytoplasmic motif. (**C**) Schematic comparison of the two proteins showing conserved domain structure and percent identity between individual sequence elements.

### Expression of TcFra isoforms in *Drosophila* using GAL4/UAS

As a first step in comparing the functions of DmFra and TcFra, we made transgenic *Drosophila* lines to express both TcFra isoforms in *Drosophila* embryonic neurons using the GAL4/UAS system. We crossed *UAS-TcFraS* and *UAS-TcFraL* transgenes to *elav-GAL4* (which is expressed broadly in all neurons) and *ap-GAL4* (which is expressed in a subset of ipsilateral neurons which do not normally express Fra). We found that, compared to a control *UAS-DmFraS* line, both TcFra isoforms were expressed at much lower levels and were unable to promote ectopic midline crossing of the apterous neurons **(Figure S1)**. The unexpected weak expression levels of our UAS transgenes prevented us from making any conclusions about the abilities of the TcFraL and TcFraS proteins to influence midline crossing in *Drosophila*. Instead, to better compare the axon guidance activities of DmFra and TcFra in *Drosophila* neurons in the pattern and expression levels of endogenous *fra*, we turned to a CRISPR/Cas9-based gene replacement approach.

### CRISPR mediated gene replacement of *Drosophila frazzled*

We used CRISPR/Cas9-mediated homology-directed repair (HDR) to replace the *Drosophila fra* gene with cDNAs encoding HA-tagged DmFraS or TcFraS proteins **(Figure 2)** and compared the expression and function of the two proteins when expressed from the endogenous *fra* locus. We previously used this approach to successfully replace *Drosophila robo3* with *Tribolium Robo2/3* and showed that TcRobo2/3 protein was properly expressed in *Drosophila* embryonic neurons and could fully substitute for *Drosophila* Robo3 to guide longitudinal axons in the intermediate region of the embryonic nerve cord (Evans, 2017).

**Figure 2.**
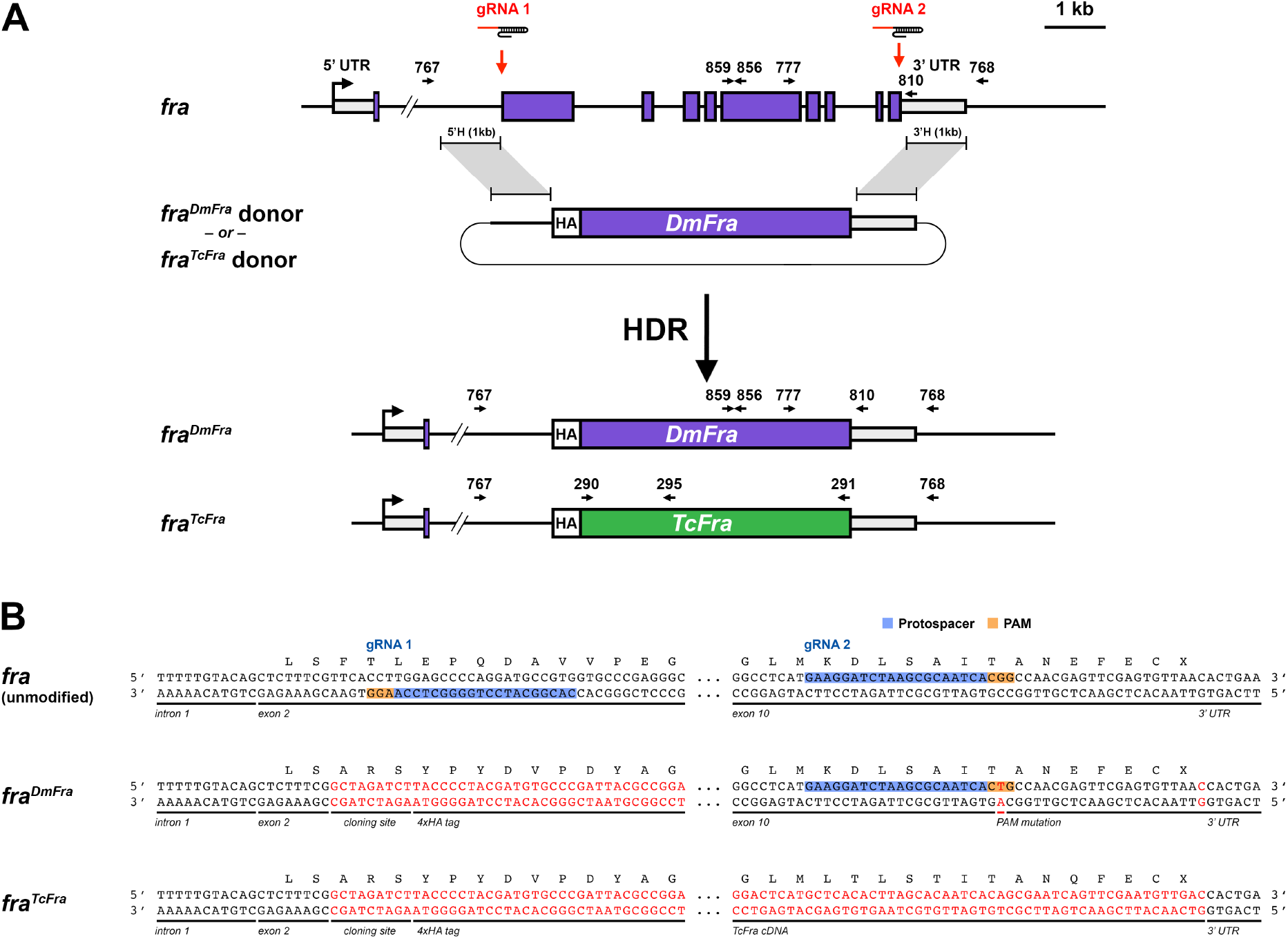
CRISPR-based gene replacement of *Drosophila fra*. (**A**) Schematic of the *fra* gene showing intron/exon structure and location of gRNA target sites, homologous donor plasmid, and the final modified *fra*^*DmFra*^ and *fra*^*TcFra*^ alleles. Endogenous *fra* coding exons are shown as purple boxes; 5′ and 3′ untranslated regions are shown as light gray boxes. The start of transcription is indicated by the bent arrow. Introns and exons are shown to scale, with the exception of the first intron, from which approximately 24 kb has been omitted. Red arrows indicate the location of upstream (gRNA 1) and downstream (gRNA 2) gRNA target sites. Gray brackets demarcate the region to be replaced by sequences from the donor plasmid. (**B**) Partial DNA sequences of the unmodified *fra* gene and the modified *fra*^*DmFra*^ and *fra*^*TcFra*^ alleles. Black letters indicate endogenous DNA sequence; red letters indicate exogenous sequence. Both DNA strands are illustrated. The gRNA protospacer and PAM sequences are indicated for both gRNAs. The first eight base pairs of *fra* exon 2 are unaltered in the CRISPR modified alleles, and the *fra* coding sequence beginning with codon F37 is replaced by the HA-tagged *DmFra* or *TcFra* cDNA sequences in *fra*^*DmFra*^ and *fra*^*TcFra*^ respectively. The endogenous *fra* transcription start site, ATG start codon, and signal peptide are retained in exon 1. The PAM and protospacer sequences for the two gRNAs are not present in the *fra*^*TcFra*^ modified allele, and both PAM sequences are mutated in the *fra*^*DmFra*^ modified allele, ensuring that the modified sequences are not cleaved by Cas9. UTR, untranslated regions; 5’H, 5’ homology region; 3’H, 3’ homology region; HA, hemagglutinin epitope tag; gRNA, guide RNA; HDR, homology directed repair; PAM, protospacer adjacent motif.

Here, we used two guide RNAs (gRNAs) to target Cas9-induced double strand breaks in exons 2 and 10 of the *Drosophila fra* gene, and provided donor plasmids carrying HA-tagged *DmFra* or *TcFra* coding sequences flanked by 1 kb homology arms as HDR repair templates.

*Drosophila* embryos expressing Cas9 in germline cells (*nos-Cas9)* (Port et al., 2014) were injected with a mix of gRNA-encoding plasmid and donor plasmid, and the progeny of these injected individuals were screened by PCR to identify HDR events. We recovered balanced stocks of the modified *fra*^*DmFra*^ and *fra*^*TcFra*^ alleles and sequenced the modified loci to confirm replacement of the endogenous *fra* locus and correct integration of the donor sequences.

Amorphic alleles of *fra* are homozygous lethal, while flies homozygous for the *fra*^*DmFra*^ or *fra*^*TcFra*^ alleles are viable and fertile, indicating that both replacements can rescue the essential functions of *fra*.

### Expression of HA-tagged DmFra and TcFra in *Drosophila* embryos

During embryonic development, *Drosophila* Fra is broadly expressed throughout the central nervous system (CNS), and Fra protein is detectable on both longitudinal and commissural axons in the ventral nerve cord, as well as motor axons that innervate muscles in the peripheral body wall (Kolodziej et al., 1996). In addition to its nervous system expression, Fra is also expressed in other parts of the developing embryo, including epidermis and midgut epithelial cells. We examined DmFra and TcFra expression in embryos homozygous for our HA-tagged *fra*^*DmFra*^ or *fra*^*TcFra*^ alleles using anti-HA antibodies.

Consistent with previous reports of endogenous Fra protein expression, we observed strong HA expression in the developing midgut, epidermis, and central nervous system **(Figure 3)** (Kolodziej et al., 1996). Importantly, we could detect axonal localization of DmFra and TcFra protein in the ventral nerve cord, indicating that both proteins are stably expressed and properly localized in *Drosophila* embryonic neurons, and that addition of the N-terminal 4xHA tag and removal of the eight intervening introns in the *fra* locus did not interfere with these aspects of protein expression or localization. We also observed similar levels of expression in *fra*^*DmFra*^ and *fra*^*TcFra*^ embryos, indicating that the *DmFra* and *TcFra* coding sequences are transcribed and translated at equivalent levels in these modified backgrounds.

**Figure 3.**
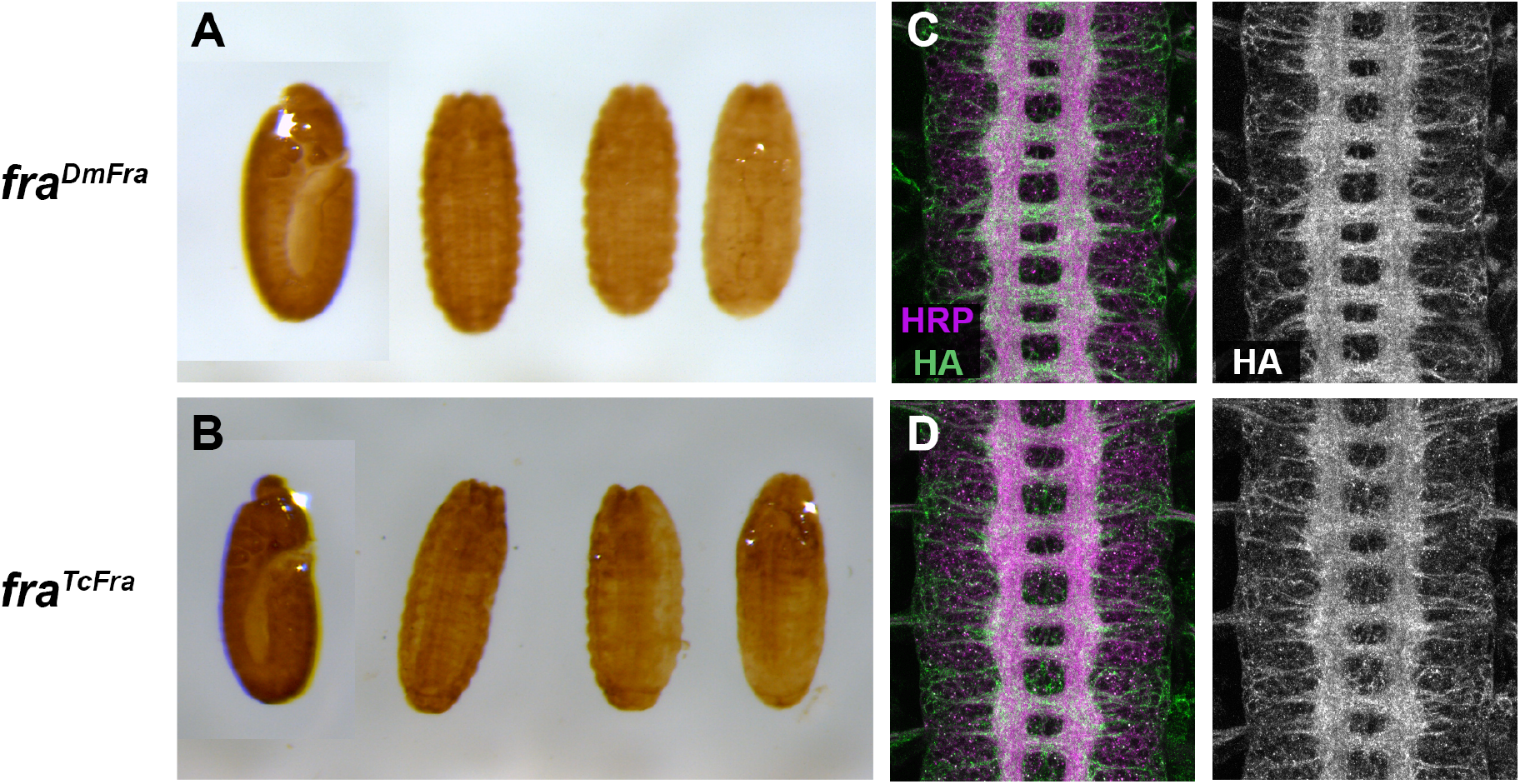
Expression of DmFra and TcFra proteins in *Drosophila* embryos. **(A**,**B)** Whole mounted *Drosophila* embryos stained with anti-HA to detect HA-tagged DmFra protein in *fra*^*DmFra*^ embryos **(A)** and TcFra protein in *fra*^*TcFra*^ embryos **(B)**. Both proteins are broadly expressed throughout the epidermis, central nervous system, and gut. **(C**,**D)** Stage 16 embryos fluorescently stained with anti-HRP (magenta; labels all axons) and anti-HA (green) antibodies. Panels at right show isolated anti-HA channels. HA-tagged DmFra **(C)** and TcFra **(D)** protein are present at high levels on longitudinal and commissural axons in the ventral nerve cord, reproducing DmFra’s endogenous expression pattern. The overall morphology of the axon scaffold appears wild type in embryos of both genotypes.

**Figure 4.**
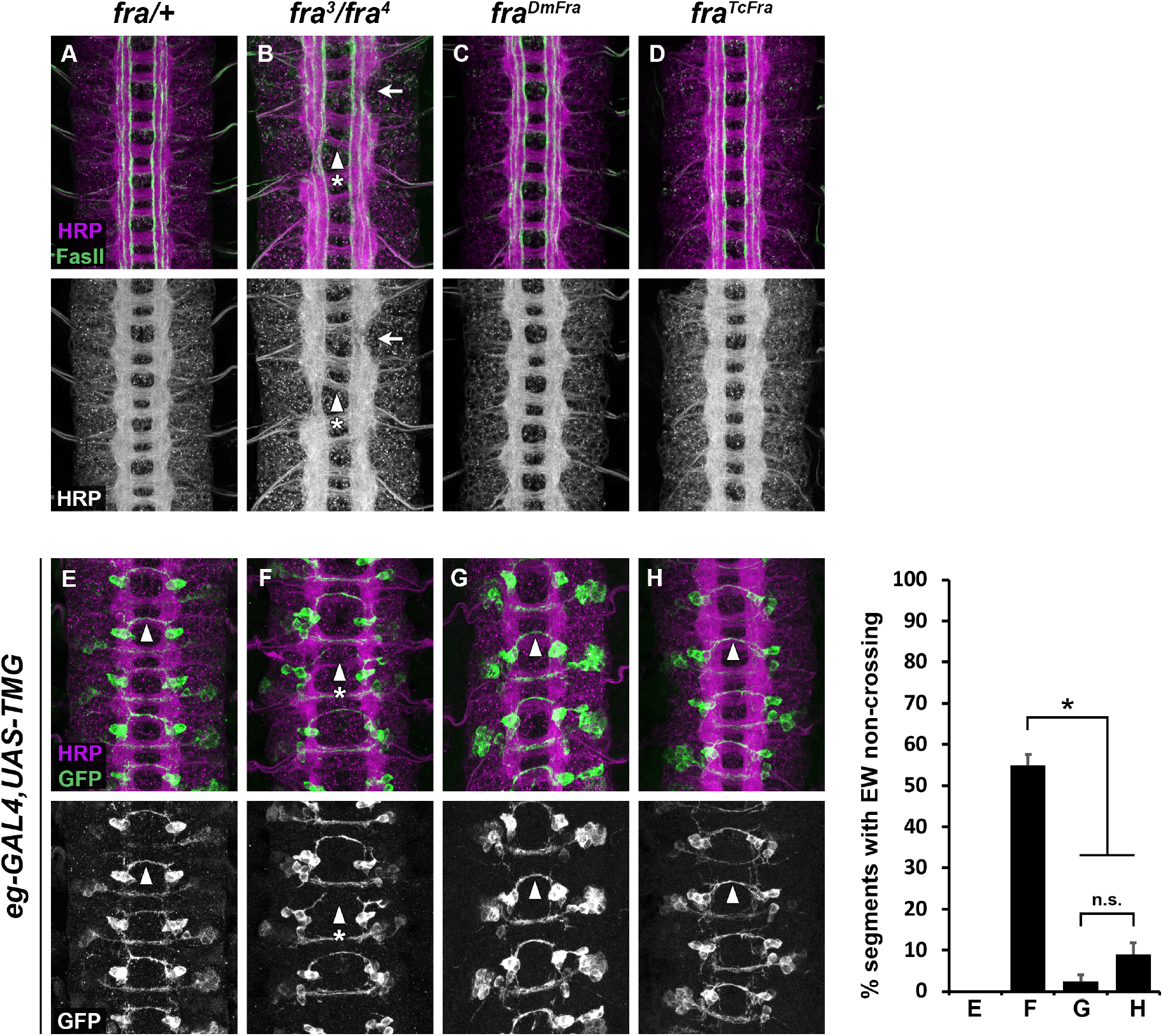
*Tribolium* Fra can substitute for *Drosophila* Fra to promote midline crossing. **(A-D)** Stage 16-17 embryos stained with anti-HRP (magenta) and anti-FasII (green; labels a subset of longitudinal axons) antibodies. In *fra/+* embryos **(A)**, the ventral nerve cord axon scaffold has a regular ladder-like structure with two commissures in each segment. In *fra* mutants, midline crossing is reduced producing thinning or absent commissures **(B, arrowhead with asterisk)**, and breaks form in the longitudinal connectives **(arrow)**. The ventral nerve cords of *fra*^*DmFra*^ **(C)** and *fra*^*TcFra*^ **(D)** embryos are indistinguishable from wild type. **(E-H)** Stage 15-16 embryos carrying *eg-GAL4* and *UAS-TauMycGFP* (*TMG*), stained with anti-HRP (magenta) and anti-GFP (green) to label the EG and EW commissural neurons. In *fra/+* embryos **(E)**, the EW axons cross the midline in the posterior commissure in every segment of the ventral nerve cord **(arrowhead). (F)** In *fra* mutant embryos (*fra*^*3*^*/fra*^*4*^), EW axons fail to cross the midline in 58% of segments **(arrowhead with asterisk)**. EW midline crossing is nearly completely rescued in *fra*^*DmFra*^ **(G)** and *fra*^*TcFra*^ **(H)** embryos. Bar graph quantifies EW crossing defects in the genotypes shown in (E-H). Error bars show s.e.m. Percent defects for the two modified alleles were compared to *fra* mutants and each other by two-tailed Student’s t-test with a Bonferroni correction for multiple comparisons (*p<10e-8; n.s., not significant). Twenty embryos were scored for each genotype.

To more closely examine the expression of DmFra and TcFra in the ventral nerve cord, we performed confocal microscopy on embryos stained with anti-HA in combination with an antibody against horseradish peroxidase (anti-HRP), which cross-reacts with a pan-neural epitope in *Drosophila* embryos and thus labels all of the axons in the ventral nerve cord (Jan and Jan, 1982; Snow et al., 1987). In homozygous *fra*^*DmFra*^ embryos, HA-tagged DmFra protein was strongly expressed in the ventral nerve cord, and primarily localized to neuronal axons.

DmFra protein was present at similar levels on longitudinal and commissural axon tracts, again consistent with previous reports of endogenous Fra protein expression. We observed equivalent expression levels and protein localization of TcFra protein in homozygous *fra*^*TcFra*^ embryos, confirming that the TcFra protein is properly translated and stably localized in *Drosophila* neurons.

### *Tribolium* Fra can substitute for *Drosophila* Fra to promote midline crossing of axons in the ventral nerve cord

We next characterized *fra*-dependent axon guidance decisions in our modified backgrounds to determine if TcFra could substitute for DmFra to attract axons toward and across the midline. We again used anti-HRP to label the entire axon scaffold, and combined this with an antibody against Fasciclin II (anti-FasII), which labels a subset of longitudinal axon pathways in the ventral nerve cord (Grenningloh et al., 1991; Harrelson and Goodman, 1988). In wild type embryos or *fra/+* heterozygotes, the two parallel longitudinal connectives on either side of the midline and two commissures in each segment form a regular ladder-like axon scaffold, with FasII-positive longitudinal tracts located in three discrete regions of the neuropile. In *fra* mutants, defects are observed in both commissural and longitudinal axon guidance: midline crossing is reduced, resulting in thin or absent commissures; and breaks appear in the longitudinal connectives along with disorganization of FasII-positive axons, reflecting defects in longitudinal axon growth and/or guidance (Kolodziej et al., 1996). The overall appearance of the axon scaffolds in both *fra*^*DmFra*^ and *fra*^*TcFra*^ embryos were indistinguishable from wild type, with no detectable thinning of commissures, longitudinal breaks, or defects in FasII pathway formation that would indicate a reduction in *fra* function in these embryos.

To quantify *fra-*dependent midline crossing in our modified embryos, we labeled a subset of commissural axons using *eg-GAL4* and *UAS-TauMycGFP (UAS-TMG)* transgenes in homozygous *fra*^*DmFra*^ and *fra*^*TcFra*^ embryos and scored midline crossing of the EW commissural neurons compared to wild type and *fra* mutant control embryos. In *fra* mutant embryos, EW axons fail to cross the midline in 58% of segments, compared to 0% in *fra/+* heterozygous siblings. EW midline crossing defects are significantly rescued in homozygous *fra*^*DmFra*^ embryos (2.5% EW non-crossing) and homozygous *fra*^*TcFra*^ embryos (8.9% EW non-crossing). The degree of rescue we observed in *fra*^*DmFra*^ versus *fra*^*TcFra*^ embryos was not statistically different (p=0.06 by student’s t-test), indicating that TcFra can function as effectively as DmFra to promote midline crossing of EW axons in the *Drosophila* embryonic CNS. Together, these results demonstrate that the *Tribolium* Frazzled receptor can substitute for its *Drosophila* ortholog to properly regulate multiple aspects of axon guidance during development of the *Drosophila* embryonic CNS.

## Discussion

In this paper, we have used a CRISPR/Cas9-mediated gene replacement approach to compare the axon guidance activities of the *Tribolium* Frazzled (TcFra) receptor with its *Drosophila* ortholog (DmFra). We replaced the endogenous *frazzled (fra)* gene in *Drosophila* with HA-tagged cDNAs encoding DmFra or TcFra, and examined protein expression and localization for each engineered allele. We examined midline crossing and longitudinal axon guidance in both modified backgrounds, compared to control wild type and *fra* mutant animals. We find that TcFra protein is present at equivalent levels and exhibits the same axonal localization pattern as DmFra when expressed from the endogenous *fra* locus, and that axon guidance in the embryonic CNS occurs normally in *fra*^*DmFra*^ and *fra*^*TcFra*^ homozygous embryos. Together, these results demonstrate that TcFra can substitute for DmFra to regulate axon guidance during embryonic development, and support the hypothesis that *Drosophila* and *Tribolium* Fra can promote midline crossing via equivalent mechanisms.

We have previously used a similar gene replacement approach to create engineered alleles of *Drosophila robo1, robo2*, and *robo3*, in which the majority of coding exons and their intervening introns are replaced with epitope-tagged cDNAs encoding wild type or modified versions of the genes themselves, or orthologs from other species (Agcaoili and Evans, 2024; Carranza et al., 2023; Evans, 2017; Howard et al., 2021). In each case, as with our experiments with *fra* reported here, neither adding an N-terminal 4xHA epitope tag nor deleting intervening introns had any detectable effect on the expression or functions of the targeted genes. One caveat to this is that we have not closely examined other aspects of expression or function outside of the embryonic CNS. For example, *fra* is expressed in the adult ovary and is necessary for female fertility (Russell et al., 2021). Although homozygous *fra*^*DmFra*^ and *fra*^*TcFra*^ adults are viable and fertile, suggesting that there are no serious defects in ovarian expression or function of *fra*. we have not directly assayed DmFra or TcFra expression in ovaries of flies carrying our engineered alleles. It is possible that one or more of the introns deleted in our gene replacement alleles might be required for expression in the ovary or other tissue(s).

In addition to signaling Netrin-dependent midline attraction, *Drosophila* Fra also activates transcription of *comm*, a negative regulator of Slit-Robo signaling, via nuclear translocation of the Fra intracellular domain (ICD) in response to an unknown signal (Neuhaus-Follini and Bashaw, 2015; Yang et al., 2009; Zang et al., 2022). *comm* does not appear to be conserved in *Tribolium*, but TcFra may regulate the transcription of other genes in *Tribolium*, such that it retains the ability to regulate *comm* when expressed in *Drosophila* neurons even though *comm* may not be among its usual target genes. The fact that we see no detectable midline crossing defects in *fra*^*TcFra*^ homozygous embryos suggests that TcFra must be able to activate *comm* transcription in *Drosophila* neurons, as we would predict that some midline crossing defects would remain if TcFra were only able to rescue Netrin-dependent attraction and not Netrin-independent activation of *comm*. Nonetheless, it will be necessary to directly assay *comm* transcription levels in *fra*^*TcFra*^ embryos to answer this definitively.

To fully understand the role(s) of TcFra in midline axon guidance in *Tribolium* will require loss of function analysis of *TcFra* in *Tribolium* embryos. Will *TcFra*-deficient beetle embryos exhibit a commissureless phenotype, like *Aedes fra* knockdown embryos (Clemons et al., 2011), or will the midline crossing defects be more mild, as in *Drosophila?* The former would indicate that Fra plays a more essential role in midline crossing in beetles than in flies. If the latter, do the same factors that promote Fra-independent midline crossing in *Drosophila* perform the same function(s) in *Tribolium?* Would knocking out *TcRobo2/3* enhance the *TcFra* phenotype, similar to *Drosophila fra,robo2* double mutants, indicating that *Drosophila* Robo2’s pro-crossing role is conserved in *Tribolium* Robo2/3? Our initial attempts at knocking down *TcFra* via parental RNAi, which we previously used effectively to knock down *TcSlit, TcRobo1*, and *TcRobo2/3* (Evans and Bashaw, 2012), resulted in female sterility (L. Terry and T.A.E., unpublished). This may reflect evolutionary conservation of *fra*’s role in oogenesis (Russell et al., 2021). As an alternative to RNAi knockdown, CRISPR-based targeted mutation could be used to investigate the functions of *TcNet* and *TcFra* in beetle embryonic development (Gilles et al., 2015).

## Materials and methods

### Molecular biology

#### Cloning of *Tribolium frazzled*

The *Tribolium frazzled* ortholog *(TcFra)* was identified by searching the *Tribolium* genome sequence by tblastn using the *Drosophila* Fra protein as a query sequence. Forward and reverse PCR primers were designed to match the predicted exons encoding the Ig1 domain and stop codon, respectively. Total RNA was extracted from 0 to 6 day old (at 25ºC) *Tribolium* embryos using TRIzol (Invitrogen) and cDNA was synthesized using Phusion RT-PCR Kit (Thermo Scientific). Partial *TcFra* coding sequences (beginning with the first immunoglobulin-like [Ig1] domain) were amplified by PCR from cDNA using primers [367-368], digested with XbaI, and ligated into an NheI/XbaI-digested p10UASTattB vector including heterologous 5′ UTR and signal sequences (derived from the *Drosophila wingless* gene) and an N-terminal 3xHA tag. Five clones were sequenced and two alternatively spliced isoforms were identified, designated *TcFraL* (long isoform; 4,359 bp from Ig1 to stop codon) and *TcFraS* (short isoform; 4,026 bp from Ig1 to stop codon). Nucleotide sequences derived from these cDNA clones were deposited in the GenBank database: accession numbers PP851978 *(TcFraL)* and PP851977 *(TcFraS)*.

#### Construction of *fra*^*TcFra*^ and *fra*^*DmFra*^ donor plasmids

The initial *fra* CRISPR donor backbone plasmid was assembled from three PCR fragments via Gibson assembly (New England Biolabs #E2611). The three fragments were derived from pIDTSmart (plasmid backbone; primers 611-612) and the wild-type *Drosophila fra* genomic locus (5′ and 3′ homology regions; primers 607-608 and 609-610). An N-terminal 4xHA epitope tag was added by ligating annealed oligos 619-620 into the NheI site located between the 5’ and 3’ homology regions, destroying the XbaI/NheI site upstream of the 4xHA sequence and leaving an intact downstream NheI site. The *Drosophila fra (DmFraS;* primers 743-744) and *Tribolium fra (TcFraS;* primers 745-746) coding sequences were amplified by PCR and cloned into the NheI-digested 4xHA backbone via Gibson assembly. The entire donor regions including coding sequences and flanking homology regions were sequenced prior to injection. Modified *fra* HDR alleles include the following amino acid residues after the N-terminal 4 × HA epitope tag: *fra*^*DmFra*^ (DmFra Q33-C1375 relative to Genbank reference sequence AAM68622); *fra*^*TcFra*^ (TcFra H1-C1341 relative to Genbank partial reference sequence XBG72028).

#### Construction of *Drosophila fra* gRNA plasmid

*Drosophila fra* gRNA sequences were cloned into the tandem expression vector pCFD4 (Port et al., 2014) via PCR using primers 617 and 618 followed by Gibson assembly using the PCR product and BbsI-digested pCFD4 backbone. For gRNA 1, an additional G nucleotide was added to facilitate transcription from the U6:1 promoter.

### Genetics

#### *Drosophila* strains

The following *Drosophila* strains, transgenes, and mutant alleles were used: *fra*^*3*^ and *fra*^*4*^ (Kolodziej et al., 1996), *fra*^*DmFra*^ and *fra*^*TcFra*^ (this study), *eg*^*Mz360*^ (*eg-GAL4*) (Dittrich et al., 1997), *ap*^*md544*^ (*ap-GAL4*) (Calleja et al., 1996; O’Keefe et al., 1998), *P{w*^*+mC*^ *= Gal4-elav*.*L}3* (*elav-GAL4*) (Ogienko et al., 2020), *P{10UAS-HA-TcFraS}86Fb (UAS-TcFraS)* and *P{10UAS-HA-TcFraL}86Fb (UAS-TcFraL)* (this study), *P{10UAS-HA-Fra}86Fb (UAS-DmFra)* (Neuhaus-Follini and Bashaw, 2015), *P{UAS-TauMycGFP}II, P{UAS-TauMycGFP}III, w*^*1118*^; *sna*^*Sco*^*/CyO,P{en1}wg*^*en11*^ *(Sco/CyOwg)*, and *y*^*1*^ *M{w[+mC] = nos-Cas9*.*P}ZH-2A w* (nos-Cas9*.*P)* (Port et al., 2014). All crosses were carried out at 25 °C.

#### Generation and recovery of CRISPR-modified alleles

The *fra* gRNA plasmid was co-injected with the *fra*^*DmFra*^ or *fra*^*TcFra*^ donor plasmids into *nos-Cas9*.*P* embryos (Bloomington *Drosophila* Stock Center stock #54591) (Port et al., 2014) by BestGene Inc. (Chino Hills, CA). Injected individuals (G0) were crossed as adults to *Sco/CyOwg*. Founders (G0 flies producing F1 progeny carrying modified alleles) were identified by testing two pools of three F1 females per G0 cross by genomic PCR. From each identified founder, 5–10 F1 males were then crossed individually to *Sco/CyOwg* virgin females. After 3 days, the F1 males were removed from the crosses and tested by PCR with the same set of primers to determine if they carried the modified allele. F2 flies from positive F1 crosses were used to generate balanced stocks, and the modified alleles were fully sequenced by amplifying the modified locus from genomic DNA, then sequencing the PCR product after cloning via CloneJET PCR cloning kit (Thermo Scientific).

#### Immunofluorescence and imaging

*Drosophila* embryo collection, fixation, and antibody staining were carried out as previously described (Kidd and Evans, 2023a; Patel, 1994). The following antibodies were used: mouse anti-Fasciclin II (Developmental Studies Hybridoma Bank [DSHB] #1D4, 1:100), mouse anti-βgal (DSHB #40-1a, 1:150), mouse anti-HA (BioLegend #901503, 1:1000), rabbit anti-GFP (Invitrogen #A11122, 1:1000), Alexa 488-conjugated goat Anti-HRP (Jackson Immunoresearch #123-545-021, 1:500), Alexa 647-conjugated goat anti-HRP (Jackson #123-605-021, 1:100), Cy3-conjugated goat anti-mouse (Jackson #115-165-003, 1:1000), Alexa 488-conjugated goat anti-rabbit (Jackson #111-545-003, 1:500), and HRP-conjugated goat anti-mouse (Jackson ImmunoResearch #115-035-003, 1:500). Embryos were genotyped using balancer chromosomes carrying *lacZ* markers, or by the presence of labeled transgenes. Embryos stained with HRP-conjugated antibodies were developed by incubation with Stable Diaminobenzidine (DAB) solution (Invitrogen #750118) according to the manufacturer’s instructions. Stained embryos were stored in 70% glycerol/PBS at room temperature. Ventral nerve cords from embryos of the desired genotype and developmental stage were dissected and mounted in 70% glycerol/PBS (Kidd and Evans, 2023b). Fluorescent confocal stacks were collected using a Leica SP5 confocal microscope and processed by Fiji/ImageJ (Schindelin et al., 2012) and Adobe Photoshop software.

## Acknowledgments

We thank Alexandra Neuhaus-Follini and Greg Bashaw for providing *UAS-DmFra* flies and plasmid DNA. Stocks obtained from the Bloomington Drosophila Stock Center [National Institutes of Health (NIH) grant P40 OD-018537] were used in this study. Monoclonal antibodies were obtained from the Developmental Studies Hybridoma Bank, created by the Eunice Kennedy Shriver National Institute of Child Health and Human Development of the NIH and maintained at The Department of Biology, University of Iowa, Iowa City, IA 52242. This work was supported by NIH grant R15 NS-098406 (T.A.E.).

---

**Figure S1.**
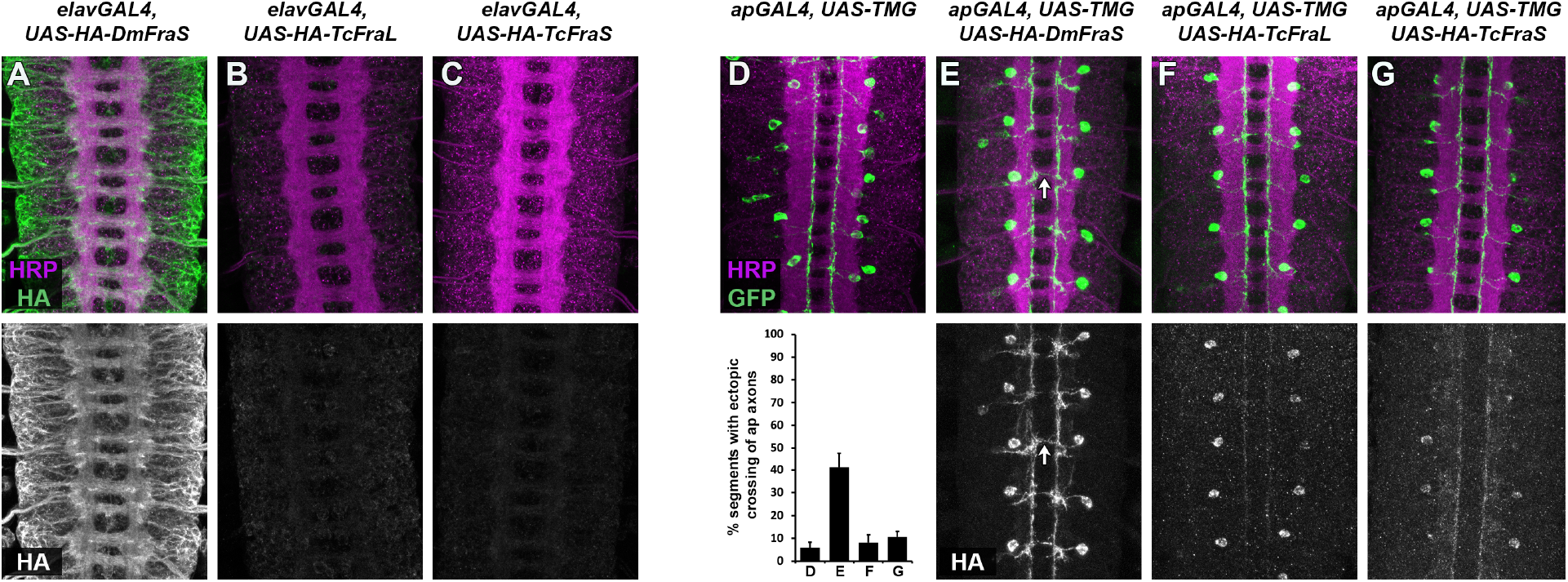
Expression of TcFra isoforms in *Drosophila* using GAL4/UAS. **(A-C)** Stage 16 embryos carrying *elav-GAL4* along with HA-tagged *UAS-DmFra* **(A)** or *UAS-TcFra* **(B**,**C)** transgenes, stained with anti-HRP (magenta) and anti-HA (green). Isolated anti-HA channels are shown below in grayscale. HA-tagged DmFraS protein is detectable at high levels in axons and cell bodies of ventral nerve cord neurons when pan-neurally misexpressed with *elav-GAL4* **(A)**. The two *UAS-TcFra* transgenes are expressed at much lower levels when combined with the same *elav-GAL4* driver **(B**,**C).** **(D-G)** Stage 16 embryos carrying *ap-GAL4* and *UAS-TauMycGFP (UAS-TMG)* transgenes alone **(D)** or in combination with HA-tagged *UAS-DmFra* **(E)** or *UAS-TcFra* **(F**,**G)** transgenes, stained with anti-HRP (magenta) and anti-GFP (green). Isolated anti-HA channels from the same embryos are shown below in grayscale. TauMycGFP expression driven by *ap-GAL4* labels around three ipsilateral neurons per abdominal hemisegment, and their axons rarely cross the midline **(D)**. Forcing the apterous neurons to express DmFra causes their axons to ectopically cross the midline in around 40% of segments **(E, arrow)**. The two *UAS-TcFra* transgenes are again expressed at much lower levels when combined with the same *ap-GAL4* driver, and they are unable to induce ectopic midline crossing of the apterous axons **(F**,**G)**. Bar graph quantifies ectopic apterous axon crossing in the genotypes shown in **(D-G)**. Error bars show s.e.m; *n* = 20-23 embryos per genotype.

